# An empirical evaluation of multivariate lesion behaviour mapping using support vector regression

**DOI:** 10.1101/446153

**Authors:** Christoph Sperber, Daniel Wiesen, Hans-Otto Karnath

## Abstract

Multivariate lesion behaviour mapping based on machine learning algorithms has recently been suggested to complement the methods of anatomo-behavioural approaches in cognitive neuroscience. Several studies applied and validated support vector regression-based lesion symptom mapping (SVR-LSM) to map anatomo-behavioural relations. However, this promising method, as well as the multivariate approach per se, still bears many open questions. By using large lesion samples in three simulation experiments, the present study empirically tested the validity of several methodological aspects. We found that i) correction for multiple comparisons is required in the current implementation of SVR-LSM, ii) that sample sizes of at least 100 to 120 subjects are required to optimally model voxel-wise lesion location in SVR-LSM, and iii) that SVR-LSM is susceptible to misplacement of statistical topographies along the brain’s vasculature to a similar extent as mass-univariate analyses.

## 1 Introduction

Studies on patients with focal brain lesions are a main source of our knowledge on the anatomo-behavioural architecture of the brain (Rorden & Karnath, 2004). For statistical analysis of lesion anatomy, different approaches of voxel-based lesion behaviour mapping (VLBM) have been implemented (Bates et al., 2003; Rorden et al., 2007). The main idea behind VLBM is to test each brain voxel individually if damage to the voxel is associated with a certain behavioural measure. As this is performed by computing a univariate test at each voxel, VLBM has also been termed a ‘mass-univariate’ approach.

While over a hundred studies have utilised VLBM so far (see Karnath & Rennig, 2017), the mass-univariate approach — like all modern neuroimaging techniques — has limitations (Karnath et al., 2018). The central problem of mass-univariate analyses is the fact that multiple univariate tests are per se independent. The assumption of independence, however, does not appear to be appropriate in investigating lesion-deficit relations. First, brain functions are not organised in single voxels, but in larger anatomical modules or networks. Second, stroke lesions do not damage the brain in a voxel-wise, independent manner, but — due to vasculature — systematically with typical patterns of collateral damage.

Previous studies have addressed these issues empirically. Simulation studies have shown that VLBM might fail to identify cognitive modules organised in a network (Mah et al., 2014; Zhang et al., 2014; Pustina et al., 2018). The underlying problem has been termed the ‘partial injury problem’ (Kinkingnéhun et al., 2007; Rorden et al., 2009). This problem appears in VLBM if a cognitive module is only partially injured by lesions, e.g. if only parts of a brain network are damaged. Statistical power can then be reduced, and VLBM might fail to identify the cognitive module in parts or in whole (see Fig. 1 for an illustration).

**Figure 1:**
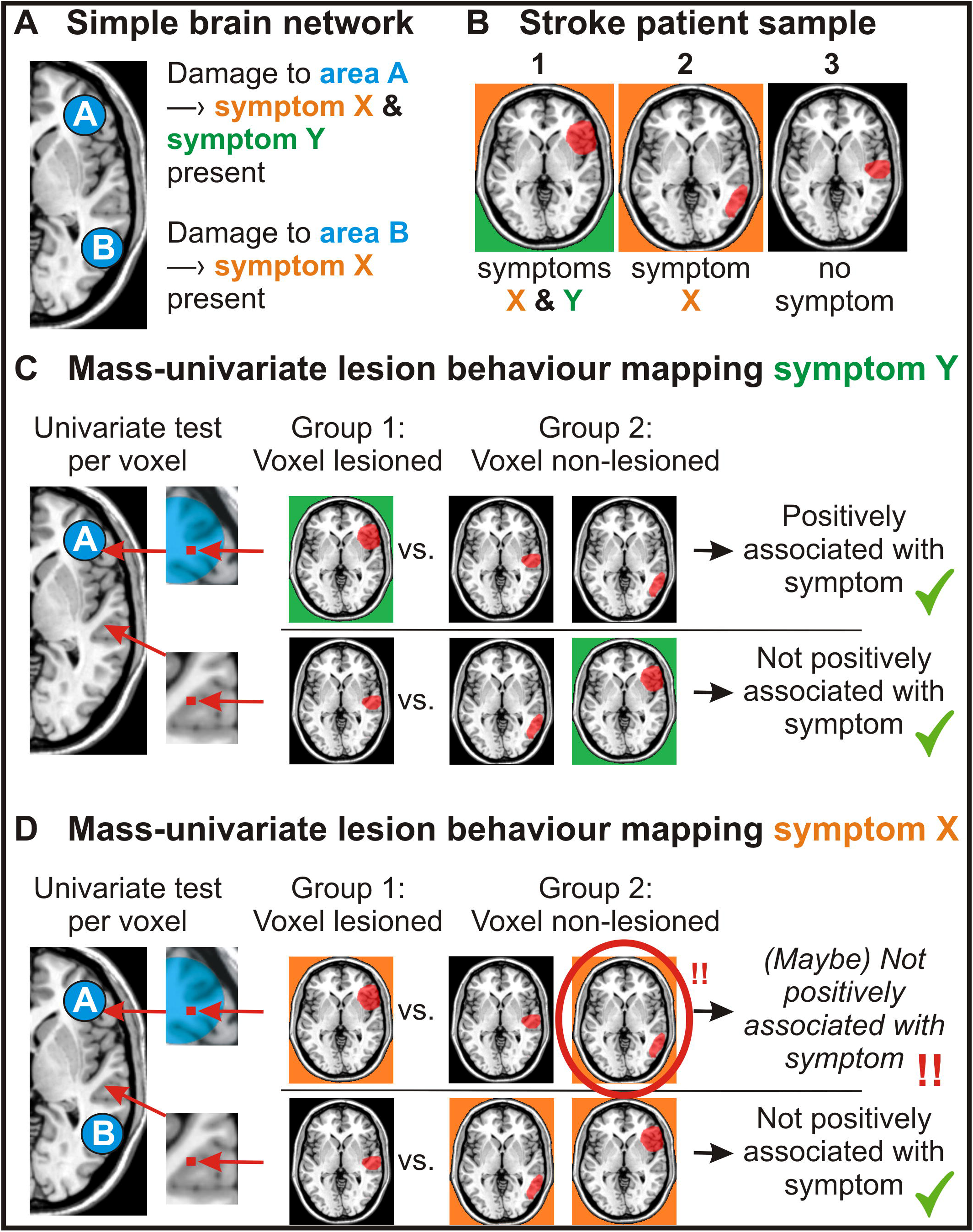
The ‘partial injury problem’. Illustration of the ‘partial injury problem’ in mass-univariate lesion behaviour mapping. **(A)** A simple fictional brain network consisting of two nodes. Damage to either node causes the same symptom X, while only damage to area A induces symptom Y. **(B)** A stroke sample of three patients. Note that the neural correlates of symptom X are *partially injured* in patients 1&2 **(C)** Mass-univariate lesion behaviour mapping of symptom Y shown for two example voxels. Following the mass-univariate VLBM approach, for each voxel patients with damage to this voxel (Group 1) are statistically tested against patients without damage to this voxel (Group 2). Voxels are considered to be associated with a symptom if Group 1 is significantly associated with a more severe symptom. For symptom Y, where damage to the brain module is either complete or not present at all, VLBM results will be correct. **(D)** Mass-univariate lesion behaviour mapping of symptom X. Here, statistical power is decreased because patients with present symptoms due to lesions in other voxels (red circle) serve as counter-examples. This can reduce the ability of mass-univariate analyses to correctly identify brain networks or large neuroanatomical modules in a whole.

Other simulation studies have identified a misplacement of VLBM results towards the centres of the arterial territories (Inoue et al., 2014; Mah et al., 2014). This bias originates from systematic collateral damage between voxels, i.e. from high correlation/dependence of lesion status between voxels (Sperber & Karnath, 2017).

To overcome these issues of mass-univariate analyses, multivariate lesion behaviour mapping (MLBM) has been suggested (Smith et al., 2013; Mah et al., 2014; Zhang et al., 2014; review in Karnath et al., 2018). In MLBM, behaviour is modelled in one single model based on the lesion status of multiple voxels or regions of interest. This can be achieved by using machine learning algorithms such as support vector machines, including support vector regression (SVR; Vapnik, 1995). Several simulation studies have shown that MLBM is indeed superior to VLBM in detecting brain networks (Mah et al., 2014; Zhang et al., 2014; Pustina et al., 2018).

While it seems that MLBM is able to overcome the partial injury problem, it has not been investigated yet, how much MLBM is susceptible to misplacement due to collateral damage between voxels. A recent study found that misplacement in multivariate analyses is low compared to a non-parametric VLBM approach (Pustina et al., 2018). However, it was not investigated if the remaining misplacement occurs spatially random or if it still occurs systematically along the brain’s vasculature.

Another open question concerns the sample sizes required for MLBM. Multivariate models naturally contain a large number of variables. Therefore, MLBM might require much larger sample sizes for parameter estimation than VLBM. A recent study investigated the performance of VLBM and MLBM at different sample sizes, and MLBM was found to be equal or even superior to VLBM also with smaller sample sizes (Pustina et al., 2018). But still, it is not known how many subjects are required to obtain a ‘good’ multivariate model.

A third issue of discussion relates to the way statistical inference is computed in MLBM. Until now, the most often used multivariate method is based on support vector regression (SVR-LSM; Zhang et al., 2014, Mirman et al., 2015; Fama et al., 2017; Griffis et al., 2017; Ghaleh et al., 2018; Wiesen et al., submitted). SVR-LSM has several advantages over other multivariate methods: the analysis can be performed voxel-wise on a whole brain-level, and continuous behavioural variables can be modelled. The groundwork of these advantages was a novel way to determine voxel-wise statistical significance. In short, SVR generates a β-parameter for each input variable (i.e. for each voxel in SVR-LSM). Contribution of β-parameters to the multivariate model is then statistically tested by permutation testing (Zhang et al., 2014). However, there is a dissent on the practically highly relevant question if correction for multiple comparisons as an additional step in SVR-LSM is required. Fama et al. (2017) argued that “because SVR-LSM considers all voxels simultaneously in a single regression model, correction for multiple comparisons is not required”. Further, Gaonkar et al., (2013) postulated that the “interdependence [of the parameters in a SVR model] has the potential to alleviate multiple comparisons problems” when used to assess voxel-wise significance in multivariate imaging analyses. On the other hand, other studies performed SVR-LSM but corrected for multiple comparisons (Griffis et al., 2017; Ghaleh et al., 2018).

The present paper aimed to scrutinise the SVR-LSM method and answer the questions outlined above by empirical means. By using simulations, we investigated three questions: i) Is a correction for multiple comparisons required in SVR-LSM? ii) What sample size is required in SVR-LSM? iii) Does SVR-LSM suffer from a misplacement of results towards the centres of the brain’s vascular territories?

## 2 General Methods

Imaging data of patients with first acute unilateral right stroke admitted to the Centre of Neurology at Tübingen University Hospital were used. Only patients with a clearly demarcated, non-diffuse lesion visible in structural imaging were included. Patients or their relatives consented to the scientific use of their data. The study has been performed in accordance with the ethical standards laid down in the 1964 Declaration of Helsinki.

Structural brain images acquired as part of clinical protocols by either CT or MRI were used for lesion mapping. If both imaging modalities were available, MRI was preferred. In patients where MR scans were available, we used diffusion-weighted imaging (DWI) if the images were acquired within 48 h after stroke onset or T2-weighted fluid attenuated inversion recovery (FLAIR) images for later scans. Lesions were manually delineated on transversal slices of the individual scan using MRIcron (www.mccauslandcenter.sc.edu/mricro/mricron). Scans were then warped to 1×1×1mm MNI space (Collins et al., 1994) using SPM8 (www.fil.ion.ucl.ac.uk/spm) and Clinical Toolbox (Rorden et al., 2012).

Multivariate lesion behaviour mapping by SVR was performed using MATLAB 2017b, libSVM (Chang and Lin, 2011), and a publicly available collection of scripts for SVR-LSM from the study by Zhang et al. (2014). Lesion maps and behavioural data were processed to fit the input data structure of libSVM and an epsilon-SVR with radial basis function kernel was computed. The resulting β-parameters were then remapped into three-dimensional MNI space. Voxel-wise statistical significance level in this parameter map was determined by permutation testing. Using this approach, data are permuted several thousands of times and the resulting pseudo-behaviour data are used to generate SVR-β-maps. Finally, voxel-wise significance is determined by comparison of pseudo-behaviour β-maps and the β-map obtained from real behavioural data. In the present study 1000 permutations were used in all experiments. Zhang and co-workers have also shown that a control for the effect of lesion size is required in SVR-LSM. Therefore, the binary lesion images were first vectorised and normalised to have a unit norm, which serves as a direct total lesion volume control (dTLVC). Derivation of statistical maps via permutation testing, kernel choice, pre-processing of behavioural variables via normalisation, back-projection of data into three-dimensional space, and control of lesion size were performed with scripting provided by Zhang et al. (2014). We consistently set hyperparameters C = 30 and *γ* = 4, which have proven to perform well in a previous study (Wiesen et al., submitted). Note that hyperparameter selection via cross validation is an important step in SVR-LSM, and has potential to maximize model quality. However, its computational demands are high, and no clear criteria are available for parameter choice yet (see Zhang et al., 2014). All further analyses were performed using MATLAB and SPSS 19; all statistical tests were computed at p < .05. Voxel-based lesion behaviour mapping was performed using NiiStat (https://www.nitrc.org/plugins/mwiki/index.php/niistatiMainPage).

## 3 Experiment 1: Correction for multiple comparisons

### *3.1* Methods

Experiment 1 aimed to clarify if a correction for multiple comparisons is required in SVR-LSM as implemented by Zhang et al (2014). Therefore, we evaluated false positive rates in lesion-behaviour samples without any actual underlying positive signal. To obtain such samples, we utilised a sample of 203 right brain damaged patients with normalised lesion maps recruited in a recent study by Wiesen et al. (submitted). Wiesen and co-workers found large significant clusters to underlie spatial neglect, using SVR-LSM both with and without correction for multiple comparisons via false discovery rate (FDR). In the present study, we permuted 20 times a continuous measure of spatial neglect behaviour (Rorden & Karnath, 2010) in this sample of 203 right brain damaged patients. The resulting samples thus did not contain any true signal. Note that we also wanted to exclude the possibility that control for lesion size affects the amount of false alarms. Therefore, SVR-LSM was performed with these 20 samples both with and without correction for lesion size (see above section ‘2 Methods’: ‘direct total lesion volume control’). Only voxels damaged in at least 10 patients were included in the modelling process. Dependent variable was the rate of voxels with false positive signal at p-levels .05 and .01.

### *3.2* Results

The analyses were performed for 349512 voxels. A large amount of false positive findings was observed in all conditions. If statistical significance is determined for each voxel independently — and independent of the fact that only a single multivariate model is computed — each analysis should yield p*349512 false positive voxels. Twotailed one-sample t-tests showed that the number of false positive voxels did neither significantly differ from these expected values for statistical parameter thresholding at p < .05 (for both t-tests p > .95; see Fig. 2) nor at p < .01 (for both t-tests p > .32). Also, paired t-tests did not find any difference between false positive rates in analyses with versus analyses without control for lesion size (both p > .98). Moreover, applying FDR correction with q = .05 to the results, none of the 40 performed analyses (20 with and 20 without control for lesion size) yielded positive results. Additionally, we performed a control analysis on random uniformly distributed data instead of permuted real data. In this analysis, we again found large numbers of false positives.

**Figure 2:**
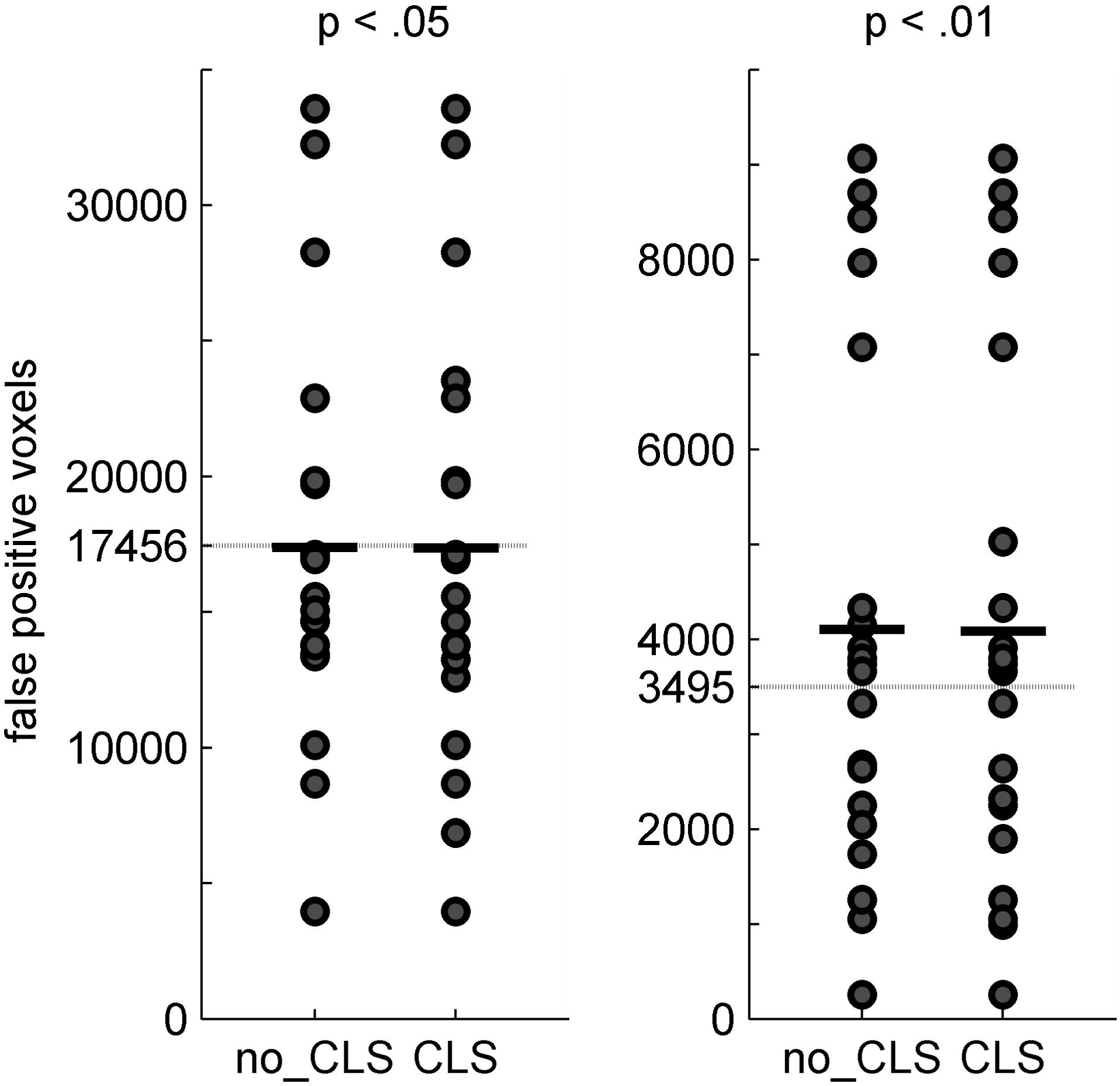
False positive rates in SVR-LSM. False positive rates of SVR-LSM in 20 simulation data sets that contain no true positive signal. Results are shown for multivariate lesion behaviour mapping with permutation-based statistical thresholding of parameter maps (see Zhang et al., 2014) at p < .05 and p < .01, and both with and without control for lesion size (CLS). Dotted lines indicate expected amount of false positives if statistical significance is indeed determined for each voxel independently; bold bars indicate mean values.

### *3.3* Discussion

At a p-level of p<.05, we found that 5% out of all tested voxels contained false positive signal (and correspondingly 1% of all voxels at p<.01). This was not affected by direct total lesion volume control. The current implementation of SVR-LSM thus poses the same challenge as VLBM: analyses find large amounts of false positive signal, and statistical maps have to be controlled for multiple comparisons. Note that this conclusion could also have been made by examining the underlying algorithms used in SVR-LSM. SVR includes a large amount of variables (here: voxels) into a single model. Generally, a comparison between two SVR models does not require a correction for multiple comparisons, although many variables are included in both models. However, permutation testing assesses statistical significance for each voxel individually. Therefore, the correction for multiple comparisons is required. From such theoretical perspective, the present empirical investigation is tautological. On the other hand, given the dissent in the field (see above section ‘1 Introduction’), the empirical approach employed in the present study provides a clear answer.

## 4 Experiment 2: Sample size required for multivariate models

### *4.1* Methods

Experiment 2 investigated what sample size is required to perform valid SVR-LSM. Six samples with simulated ‘behavioural’ data based on damage to either single or multiple areas were investigated. To obtain simulated data, 283 real normalised lesion maps of patients with right hemisphere damage were used. Most patients of this sample were also part of a recent simulation study (Sperber & Karnath, 2017). As ground truth, regions in the Automatic Anatomic Labeling (AAL) atlas (Tzourio-Mazoyer et al., 2002) were chosen. Patients’ individual ‘behavioural’ scores were then computed as a linear function of the individual lesion’s damage to the atlas region. This was a continuous score that ranged from 0 (no damage to the region) to 1 (damage to 100% of all voxels in the atlas region). For multi-region models, the score was computed based on the region with most damage. Following simulation regions were chosen: i) insula ii) middle frontal gyrus iii) inferior parietal lobule iv) inferior frontal gyrus triangular + supramarginal gyrus v) caudatum + middle temporal gyrus vi) inferior frontal gyrus triangular + supramarginal gyrus + middle temporal gyrus.

Different sample sizes up to 140 patients were investigated in steps of 20, i.e. seven different sample sizes (20, 40, 60, 80, 100, 120, 140), and it was investigated what sample size is required to obtain a valid SVR model. This leads to the non-trivial question how to assess if a multivariate model is good. The model should fit the data; however, a good fit does not imply that a model is good, as it can suffer from overfitting. Rather, a good model should also provide high generalisability. Therefore, we primarily assessed the correspondence of SVR-LSM maps, i.e. the final permutation-thresholded -maps, and the reproducibility of SVR model β-parameters (see Rasmussen et al., 2012) between distinct samples at different sample sizes. Further, we assessed the prediction accuracy via cross-validation. To investigate model performance at sample size of n lesions, 2*n lesions were randomly drawn and assigned to two exclusively disjunct samples of n lesions each. For all analyses in Experiment 2, only voxels damaged in at least 10 patients in the 2*n sample were tested. This ensured that two paired analyses were always based on the same voxels. Next, a SVR was computed to model behavioural scores based on the status of all voxels. From the SVR models and corresponding β-maps, reproducibility of β-maps and prediction accuracy were assessed. To obtain the variable ‘reproducibility of β-maps’, the correlation between β-parameters in both β-maps was computed. Note that β-weights of individual voxels only provide limited interpretability, as they only indirectly relate to the behavioural scores. Yet, reproducibility of β-weights can be interpreted in the context of generalisability with caution (as, e.g., in Rasmussen et al., 2012; Zhang et al., 2014), especially as all comparisons in the present study were based on the same behavioural variable and the same hyperparameters. Second, ‘prediction accuracy’ was assessed by applying the model obtained in the first sample for a prediction in the second sample. Then, the correlation between predicted and true behavioural values was computed. The procedure of drawing 2*n lesions and randomly assigning them to two equally large groups was repeated 50 times for each data point. The correspondence of actual SVR-LSM maps (i.e. the final β-maps obtained from β-maps via permutation testing) was assessed by also drawing 2*n lesions and computing SVR-LSM maps independently for both samples. Then, correspondence of both maps was assessed by i) comparing both FDR-corrected, thresholded maps and computing the Dice Index, which provides a measure of similarity of two binary spatial images between 0 (no spatial overlap) to 1 (maximal spatial overlap), and ii) assessing ‘reproducibility of p-maps’ by computing the correlation between p-values in both maps. In order to save computational resources, the latter procedure was repeated five times for each data point.

### *4.2* Results

Plotting the data course of the investigated variables across sample sizes (Fig. 3) revealed several noticeable features: first, the data course of the variables qualitatively differed between reproducibility of both β- and p-maps and Dice index on the one hand and prediction accuracy on the other hand. Independent t-tests revealed that reproducibility of β-maps significantly increased for all variables with each increase in sample size, except for the step from 80 to 100 patients in the simulation based on the inferior frontal gyrus. Increment-wise improvements, however, decreased rapidly with increments for larger sample sizes, and the plotted curves suggest that model performance asymptotically approaches a limiting value. Non-surprisingly, Dice indices and reproducibility of p-maps qualitatively followed a similar trend. In contrast, prediction accuracy was not significantly improved by increases in sample size above 100 subjects, while for some simulation regions prediction accuracy already peaked with sample sizes of 40 subjects. Second, standard deviations of reproducibility and prediction accuracy descriptively decreased rapidly with increasing sample sizes. Third, among simulations based on single regions (insula, middle frontal gyrus, and inferior parietal lobule), model performance differed across most data points. Independent t-tests confirmed that regarding reproducibility of β-maps, for all sample sizes, SVR performed best for simulations on the insula, and worst for simulations on the middle frontal gyrus.

**Figure 3:**
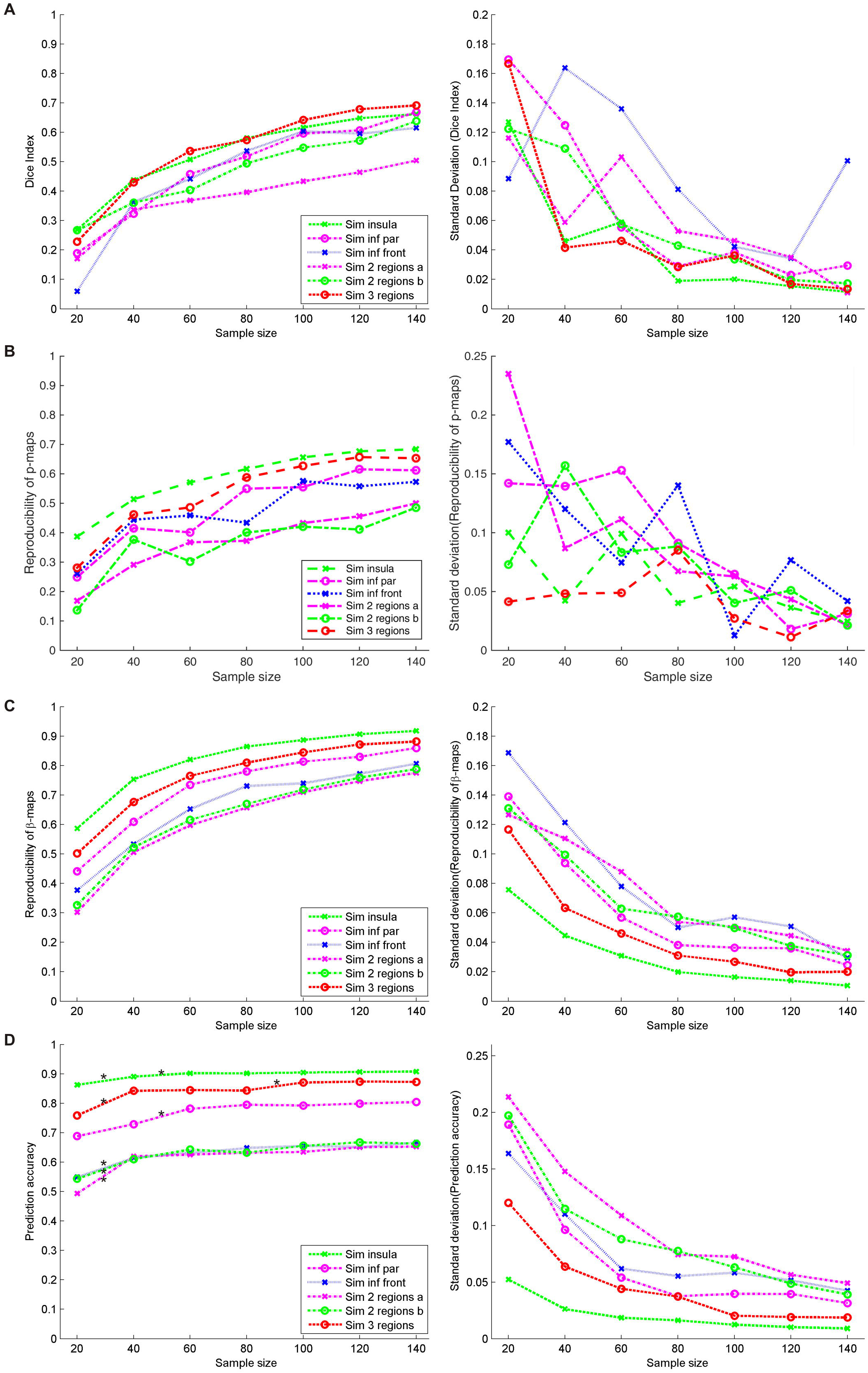
Model performance across different sample sizes. **(A)** Dice indices (left panel) and its standard deviation (right panel) at different sample sizes. Note that each data point here is based on only 5 iterations. Therefore, standard deviations are larger than for variables in panels C&D. **(B)** Reproducibility of p-maps and its standard deviation at different sample sizes. Each data point is based on 5 iterations of the experimental procedure. **(C)** Reproducibility of β-maps and its standard deviation at different sample sizes. Each data point is based on 50 iterations of the experimental procedure. All 20-subject increments except for one (see text) were connected to significant increases in reproducibility according to independent sample t-tests. **(D)** Prediction accuracy and its standard deviation at different sample sizes. Each data point is based on 50 iterations of the experimental procedure. For the left panel of Fig 3C asterisks indicate significant changes for a 20-subject increment.

### *4.3* Discussion

Improvements in the reproducibility of β-maps across all sample sizes were found, while small samples appeared to profit the most from increases in size. Increases in size based on already large samples were still significant; however, they only provided smaller benefits. A very similar trend was observed for Dice indices and reproducibility of p-maps. On the other hand, prediction accuracy was relatively stable across sample sizes and did not further improve by increases in sample size above 100 subjects. To conclude, MLBM by SVR-LSM seems to require large samples to provide a model that maximizes the use of anatomical information for parameter estimation. Optimal sample sizes appear to be larger than 140 subjects, whereas one can doubt the usefulness of increases beyond ~ 100 to 120 subjects; nominal gains in reproducibility beyond these sizes are very small. However, if SVR is not used for a parametrical mapping as in SVR-LSM, but for prediction of clinical outcome, performance already peaks with smaller sample sizes with about 40 to 100 subjects. Furthermore, with larger sample sizes standard deviations were low for both reproducibility and prediction accuracy, suggesting that model performance is quite stable for defined larger sample sizes. In other words, given a certain (larger) sample size, model parameters were equally good (or bad) across iterations.

## 5 Experiment 3: Spatial bias of statistical results

### *5.1* Methods

Experiment 3 aimed to clarify if MLBM was suffering from a misplacement of results towards the centres of the arterial territories. We thus largely copied the simulation approach to investigate misplacement of topographical results in VLBM that was used by two previous studies (Mah et al., 2014; Sperber & Karnath, 2017). Since such a simulation approach is based on highly artificial simulations, it is not perfectly transferable to real data. Yet, its simplicity fits well empirical questions that aim to assess general principles in lesion behaviour mapping (for further discussion see Mah et al., 2014). As both previous studies using this simulation approach clearly visualised misplacement on axial MNI slice z = 17, we limited the analysis to this slice. Simulations were run using a real right hemisphere lesion sample of 283 patients. In short, 464 equally distributed voxels on slice z = 17 were selected. For each voxel and patient, it was determined if the lesion includes the voxel. Contrary to previous studies, however, simulated ‘behavioural’ scores were not binary, but continuous to allow application of support vector *regression*. To do so, random normally distributed values were drawn. If a lesion included the simulation voxel, values were drawn from a distribution with μ = 1.4 and σ = 0.4, else from a distribution with μ = 0.4 and σ = 0.4. Any resulting negative scores were set to zero.

Resulting ‘behavioural’ data of 464 simulations were then used in VLBM and SVR-LSM. Only voxels damaged in at least 5 patients were considered, and all resulting topographies were corrected for multiple comparisons by FDR correction at p < .05. Misplacement was then defined as a vector ranging from the voxel the simulation was based on to the centre of mass of the thresholded, binary statistical topography. As correction for lesion size is a crucial factor in anatomical misplacement (Sperber & Karnath, 2017), VLBM and SVR-LSM were performed with two different strategies to control for lesion size. First, SVR-LSM was performed using dTLVC (Zhang et al., 2014); second, we applied an approach that controls lesion size via regression both on the behavioural variable and the anatomical data (deMarco & Turkeltaub, 2018). To perform the latter type of correction, we integrated publicly available MATLAB scripts by deMarco & Turkeltaub (https://github.com/atdemarco/svrlsmgui) into our custom-modified scripts by Zhang et al. (2014).

### *5.2* Results

For mass-univariate voxel-based lesion behaviour mapping, a misplacement of topographical results by 13.5 mm (SD = 5.8 mm; median = 13.5 mm) for analyses without control for lesion size, and by 8.7mm (SD = 4.4 mm; median = 8.2 mm) for analyses with control for lesion size by nuisance regression was found. As shown in a previous study (Sperber & Karnath, 2017), control for lesion size significantly reduced misplacement (t(887) = 13.89, p < .001). Further, a vector visualisation (Fig. 4 A and B) revealed that misplacement was systematically oriented towards the centres of the middle and posterior arterial territories. These findings thus replicated previous studies that investigated misplacement of topographical results in VLBM (Mah et al., 2014; Sperber & Karnath, 2017). Peak Z-standardised statistics in the statistical topographies were Z = 8.2 (SD = 0.7) for VLBM without control for lesion size and Z = 7.7 (SD = 0.7) for VLBM with control for lesion size.

**Figure 4:**
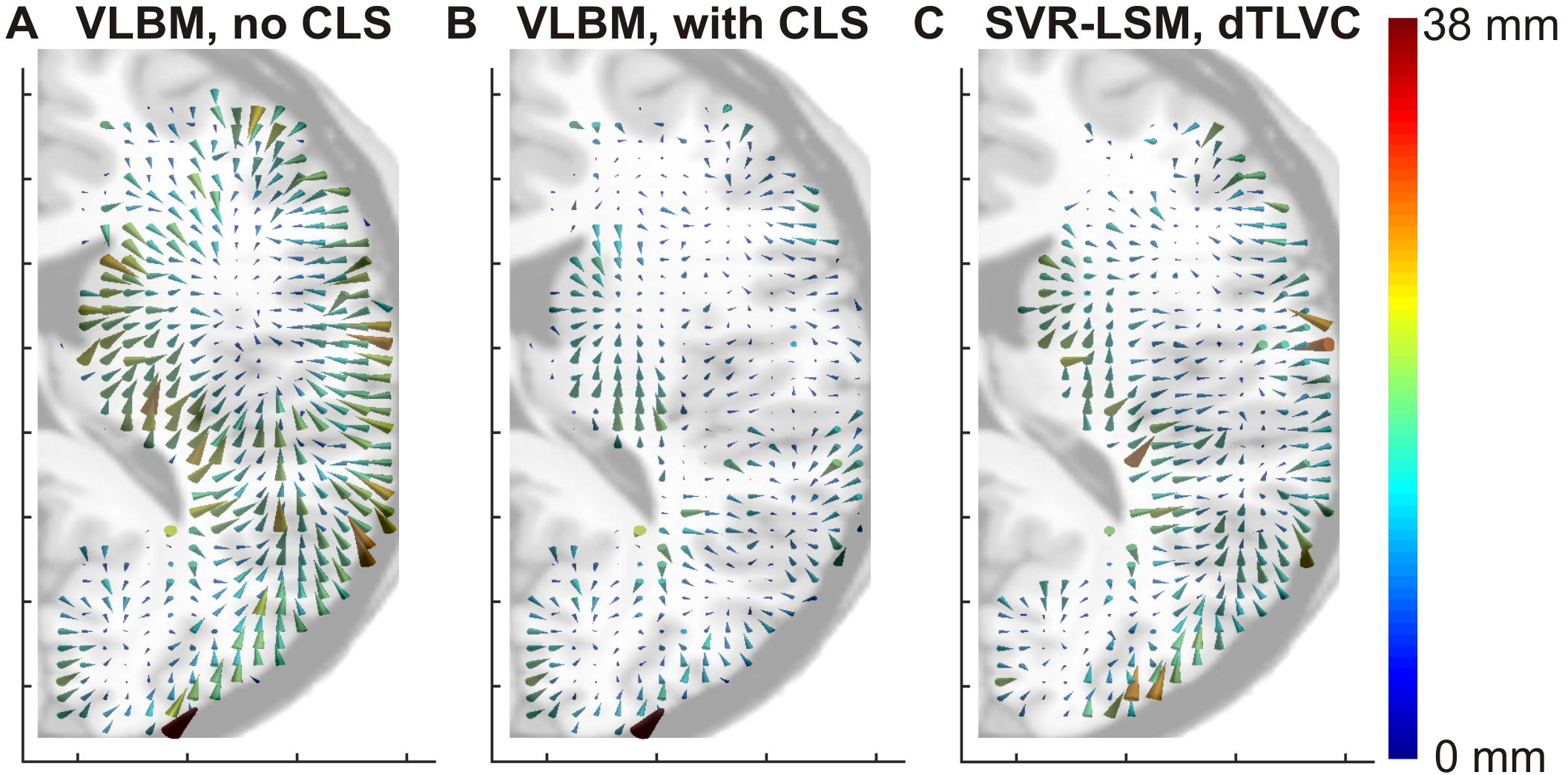
Misplacement of topographical results in VLBM and SVR-LSM. Vector maps for spatial misplacement in VLBM and SVR-LSM. Each vector shows the misplacement ranging from the ‘ground truth’ voxel to the centre of mass of the binary topographical map. All results are based on analyses controlled with FDR correction at p < .05. All analyses were limited to MNI slice z = 17. **(A)** Vector map for voxel-based lesion behaviour mapping. **(B)** Vector map for voxel-based lesion behaviour mapping with control for lesion size (CLS) via nuisance regression. **(C)** Vector map for multivariate lesion behaviour mapping using support vector regression, including a control for lesion size via dTLVC as suggested by Zhang et al. (2014).

SVR-LSM topographies with control for lesion size by dTLVC were misplaced by 11.4 mm (SD = 5.0 mm; median = 11.0 mm). This misplacement was smaller than in VLBM without control for lesion size (t(828) = 5.54, p < .001), but larger than misplacement in VLBM with control for lesion size (t(815) = 8.17, p < .001). SVR-LSM with control for lesion size by regression both on anatomy and behaviour was misplaced by 12.5 mm (SD = 7.7 mm; median = 11.2 mm), which was larger than SVR-LSM with control for lesion size by dTLVC (t(841) = 2.35; p < .05), but still smaller than uncorrected VLBM (t(913) = 2.26; p < .05). Visual inspection of the data revealed that this difference originated from few outliers; the median misplacement obtained from both methods was roughly the same.

In order to compare directionality of vectors between conditions, cosine similarity was assessed. In a comparison of two groups of vectors, average cosine similarity will be 1 if all vector pairs show the same directionality, and 0 if directionality between vector pairs is entirely random. Cosine similarity of misplacement vectors in VLBM without control for lesion size and SVR-LSM with control for lesion size by dTLVC was 0.80 (SD = 0.29), which was significantly larger than zero (t (3 78) = 54.29, p < .001). For SVR-LSM with control for lesion size by regression and VLBM without control for lesion size, cosine similarity was 0.78 (SD = 0.32), which was also larger than zero (t(378) = 48.31; p < .001. Thus, directionality of misplacement vectors in the univariate and the mass-univariate analyses were similar. Correspondingly, the vector visualisation of misplacement in SVR-LSM (Fig. 4C) also appeared to follow the vasculature.

### *5.3* Discussion

The centres of mass of statistical topographies in SVR-LSM were misplaced in a similar spatial direction as in VLBM; they were oriented towards the centres of the middle and posterior arterial territories. The magnitude of this replacement was between the magnitude of misplacement in VLBM without and VLBM with control for lesion size. Thus, SVR-LSM is not less susceptible to misplacement compared to VLBM. Different approaches to control for lesion size — by using dTLVC (Zhang et al., 2014) or via regression both on the behavioural variable and the anatomical data (deMarco & Turkeltaub, 2018) — did not eliminate the misplacement in SVR-LSM. To conclude, multivariate analysis in lesion behaviour mapping does not per se account for the complexity of lesion anatomy, and inter-voxel correlations can negatively affect results.

## 6 General Discussion

Recently, we outlined that the validity of lesion behaviour mapping methods can be tested empirically with different approaches (Sperber & Karnath, 2018). In the present study, we did so with SVR-LSM, a novel promising method with the potential to overcome some of the shortcomings of mass-univariate lesion behaviour mapping. We found that i) correction for multiple comparisons is required in SVR-LSM, ii) that sample sizes above ~ 100 to 120 subjects are required to model voxel-wise lesion location in the context of SVR-LSM, and iii) that SVR-LSM is susceptible to misplacement of statistical topographies along the vasculature in the same way as mass-univariate analyses.

Our results resolve the controversy on multiple comparisons in SVR-LSM (Zhang et al., 2014; Mirman et al., 2015; Fama et al., 2017; Griffis et al., 2017). They show that SVR-LSM requires a correction for multiple comparisons. This can be a correction by false discovery rate (FDR; Benjamini & Yekutieli, 2001) as carried out in the present study and the study by Zhang et al. (2014). However, future empirical studies are required to find the best solution to the multiple comparisons problem in SVR-LSM. For univariate analyses, several alternative solutions to the multiple comparison problem have been proposed (e.g., Nichols & Hayasaka, 2003; Rorden et al, 2007; Karnath et al., 2018; Mirman et al., 2018). Correction by FDR is easy to apply and computationally fast, and therefore well fits in a large scale simulation study. However, it has several shortcomings, e.g. if samples are of small size or only contain low signal (Karnath et al., 2018; Mirman et al., 2018). Another major limitation of FDR in SVR-LSM comes from the fact that voxel-wise p-values are obtained by permutation, which for n permutations can only have a maximal resolution of 1/n. If the number of permutations is lower than the number of statistical tests — like in probably all instances of SVR-LSM — the resolution does not suffice for mathematically correct application of FDR. It is not known yet, if this leads to marked errors in SVR-LSM, and FDR should only be used with caution and, most importantly, with larger numbers of permutations than used in the present study. The present approach simply has given way to computational limitations. A recent study introduced potential alternatives to FDR correction in SVR-LSM (DeMarco & Turkeltaub, 2018), but a consensus in the field on the optimal correction algorithm has not been found yet.

The present findings further give an answer to the question whether valid parameter estimation for multivariate analyses generally requires large data sets. Some researchers postulated that multivariate lesion behaviour mapping depends on large-scale multi-centre studies, which are able to provide such large samples (Mah et al., 2014; Xu et al., 2018). Findings in the present study partially support this assumption. Multivariate modelling based on voxel-wise lesion information in SVR consistently improved its generalisability with increases in sample size even up to 140 subjects. On the other hand, improvements beyond ~100 subjects were very small and appeared to approach a limiting value. This closely resembles findings on multivariate modelling of fMRI data (Churchill et al., 2014). The authors also observed that increases in sample size led to a plateau in regards to prediction accuracy already with small samples, while reproducibility of model parameters improved if already large samples were increased. Under the perspective of practicability, our data suggest that sample sizes of about 100 to 120 patients appear to be a good trade-off between model quality and feasibility regarding data input. It is of practical relevance to find out exactly how many patients are required to map a certain function. Cross validation, which is anyway required for hyper-parameter optimisation in SVR-LSM (see Zhang et al., 2014), can provide insights into model quality, however only smaller sub-samples of the total sample can be compared. Thus, with some limitations, cross validation could indicate if a sample was adequate in the modelling process.

However, caution should be advised if real symptoms are investigated. For real behavioural variables, using an adequate sample size for SVR-LSM would not imply that correlations close to r = 1.0 should be expected. First, anatomical information in real symptoms (compared to the present simulation samples) can vary. Critical brain regions can be organised with different complexity, spatial normalisation of brain scans is noisy, and inter-individual differences in brain anatomy exist. In other words, for real behaviours the brain-deficit relation in the normalised maps will be noisy and more complex. Second, a multitude of factors besides structural lesion information can affect post-stroke behaviour, such as age, inter-individual anatomical differences, time post stroke, or pre-stroke cognitive status (for review Price et al., 2017). Therefore, model generalisability of a SVR β-map can only be as good as behaviour can be explained by structural lesion information, e.g. voxel-wise lesion status. This makes it difficult to evaluate model performance. For example, imagine a SVR model based on a real data sample. If this model offers a cross-validation reproducibility of r = 0.4, one can hardly tell if this model already optimally includes anatomical voxel-wise information. Furthermore, differences in model performance across single-region simulations observed in the present study point at a role of lesion coverage. For simulations based on regions with higher lesion coverage (cf. Sperber & Karnath, 2016), experiment 2 has shown that models with higher lesion coverage perform better. Thus, SVR-LSM might not perform equally well for all regions, with worse performance if critical brain regions are covered by fewer lesions. Therefore, researchers that apply SVR-LSM should take care that their sample contains a considerable amount of cases in the pathological range, i.e. cases in which the critical brain region is damaged. Alternatively, investigation of behaviour that is only rarely pathological will require larger samples. Future studies on large samples of real data are required for further insights into optimal sample sizes in MLBM. Given that real data are more complex than simulated data, the present study provides a lower boundary for optimal sample sizes. Requirements for real data samples might be larger, and probably will not only be affected by sample size alone, but also by factors such as lesion distribution, or investigation of only one vs. both hemispheres at once. Such future studies could also compare different approaches of MLBM in respect to required sample sizes (e.g. Yourganov et al., 2015; Pustina et al., 2018). Experiment 2 in the present study only investigated SVR-LSM in unilateral stroke, and should not be generalised to MLBM in general. Furthermore, it should be noted that our findings do not entirely exclude studies on smaller samples, which are often found in labour-intense patient studies. However, limited generalisability of topographical results should then be communicated as a major caveat.

In the discussion of spatial misplacement inherent to mass-univariate analyses (Mah et al., 2014), it was noted that the main reason to use multivariate instead of mass-univariate analyses was the complex architecture of lesions which leads to the misplacement of statistical results (Nachev, 2015; Xu et al., 2018). In contrast, the present study suggests that multivariate lesion behaviour mapping — or at least SVR-LSM - is susceptible to misplacement of statistical topographies to the same extent as mass-univariate VLSM analyses. A simple thought experiment illustrates why these findings in fact are not surprising: Imagine a sample of 100 lesions that is used in a lesion behaviour mapping study. As commonly found in lesion samples, many voxel pairs are damaged in exactly the same lesions, so-called ‘unique patches’ (Pustina et al., 2018). In other words, there are many voxel pairs for which typical lesion anatomy leads to a perfect inter-voxel correlation of damage. Further imagine that for one of these perfectly correlated voxel pairs, one voxel belongs to a cognitive module which induces a cognitive symptom when damaged, and the other voxel does not belong to the cognitive module. In this case, we do not see any possibility that a lesion analysis - be it univariate or multivariate - could correctly identify only one voxel to belong to the cognitive module, but not the other without using any a priori information. As the present study suggests, this problem might persist for multivariate analyses even if inter-voxel correlations are not perfect, but still high.

The problem of misplacement will not be alleviated by analysing data on region level rather than on voxel level, as suggested by Nachev (2015). Lesional damage between neighbouring regions will likely correlate, and results will also be misplaced. Nevertheless, the remaining misplacement bias in SVR-LSM results — as well as in VLSM results — does not reach such levels as originally assumed in the study by Mah et al. (2014). Moreover, the quantification of misplacement as implemented in both the present and previous studies has limitations (cf. Sperber & Karnath, 2017; Pustina et al., 2018). First, a simple vector based on the centre of mass of a topographical map omits a lot of information contained in three-dimensional topographies (see Sperber & Karnath [2017] and Pustina et al. [2018] for more elaborated approaches based on more complex simulations). Second, such vectors can only point towards subcortical regions, and not towards areas outside the brain. In other words, the direction of possible biases is already predefined. This limitation might account for parts of the misplacement. However, misplacement vectors also clearly follow the arterial territories (see, Fig. 4 of the present article, or Mah et al. [2014] in comparison to e.g. Tatu et al., [2012], Neumann et al., [2016]), what indicates that lesion anatomy is a central factor in the generation of the misplacement. Another, more general limitation is the ecological validity of simulation studies. While simulations provide a powerful tool to test the validity of lesion-behaviour mapping methods (Sperber & Karnath, 2018; Xu et al., 2018), it is not known how well findings in simulations can be transferred to analyses in real data. In the present and in a previous study (Sperber & Karnath, 2017), we found peak statistics in VLBM to be very high compared to lesion studies on real data. This hints at an over-proportionally high underlying positive signal, which was present although we introduced random noise in experiment 3. The high signal, in turn, leads to more positive findings in any lesion analysis. Likely, this will induce an overestimation of misplacement in an analysis where all positive findings — except for one voxel — are false alarms. Indeed, misplacement in a simulation study has also been found to vary between p-levels of VLBM (Sperber & Karnath, 2017), with lower misplacement for more conservative p-levels, i.e. less false alarms. To conclude, as in the two previous studies using the same ‘artificial’ simulation approach (Mah et al., 2014; Sperber & Karnath, 2017), the present experiment 3 does not show that SVR-LSM topographies based on real data are misplaced by exactly ‘x’ mm (11.4mm in the present experiment), but rather that lesion anatomy generally is a biasing factor in SVR-LSM, similar as in VLBM.

The present study also bears implications for translational uses of multivariate modelling based on structural lesion data. As we discussed elsewhere (Karnath et al., 2018), these methods have potential to be used in long-term prediction of post-stroke outcome, e.g. in guiding rehabilitation measures. A recent study has shown that prediction of hemiparesis based on structural imaging can be performed with high accuracy using voxel-wise lesion information (Rondina et al., 2016). Experiment 2 in the present study contributes by showing that multivariate models maximize the use of structural lesion data already with small samples. Prediction accuracy was already relatively high even with our smallest investigated sample size of 20 patients, and did hardly improve with further increases beyond 40 to 80 subjects. However, contrary to SVR-LSM, the ultimate aim of prediction algorithms is to maximize prediction accuracy. Therefore, profound knowledge of non-anatomical variables that affect post-stroke behaviour is required (for review Price et al., 2017). Such variables can be included into SVR. However, it is not known yet if this requires larger sample sizes. Further, strategies for anatomical feature selection could improve prediction accuracies (see, e.g., Rondina et al., 2016), and other multivariate methods might be better suited to model brain-lesion relationships for prediction (Rondina et al., 2016; Hope et al., 2018). Importantly, when only prediction of behaviour is desired, both the multiple comparison problem (experiment 1) and anatomical biases (experiment 3) are not relevant.

To conclude, the present study could clarify some of the open and, in part, controversially debated questions related to SVR-LSM. Multivariate lesion behaviour mapping does not appear to resolve all methodological issues in the field of lesion-behaviour inference (yet). Nevertheless, this new and promising approach to lesion analysis enables various new insights into brain architecture. It supplements traditional mass-univariate analysis methods, in particular if larger patient samples are accessible, and might even be able to replace univariate methods entirely. In the future, MLBM methods based on other multivariate algorithms or advancements in the SVR-LSM technique might resolve present limitations of SVR-LSM.

## Acknowledgements

This work was supported by the Deutsche Forschungsgemeinschaft (KA 1258/23-1). Christoph Sperber was supported by the Friedrich Naumann Foundation. Daniel Wiesen was supported by the Luxembourg National Research Fund (FNR/11601161). The authors acknowledge support by the state of Baden-Württemberg through bwHPC.

